# Network Formation Dynamics in Thiol-ene Crosslinked Hyaluronic Acid Hydrogels: Design Principles for In Vitro Tissue Models

**DOI:** 10.64898/2026.05.17.725744

**Authors:** Kyley Burkey, Yiwen Zheng, Kinsey Drake, Ryan Brady, Cole A. DeForest, Alshakim Nelson, Aniruddh Vashisth, Jennifer Robinson

## Abstract

Hydrogels are widely used as three-dimensional cell culture systems to understand the impact of cellular mechanotransduction for tissue engineering applications. Photoinitiated thiol-ene click chemistry is a commonly utilized hydrogel crosslinking mechanism that provides spatial and temporal control over hydrogel network formation and resulting mesh size and compressive properties. Despite historically documented efficiency as step-growth reactions, these reactions do not always proceed as predicted. To understand the impact of cell confinement and microenvironmental mechanics on cellular function, thiol-ene network formation must be thoroughly characterized. To this end, the objective of this work was to investigate the crosslinking dynamics to determine hydrogel network formation as assessed via mesh size and mechanical properties using a pentenoate-functionalized hyaluronic acid thiol-ene reaction. Hydrogel parameters including polymer concentration and thiol:-ene crosslinker molar ratio were modulated (4, 6, or 8 polymer weight percent and 0.15:1, 0.5:1, or 1:1 molar ratio of thiol groups to reactive -ene groups) to tune network properties including shear storage modulus and relative mesh size. Molecular Dynamics (MD) simulations were used to simulate the thiol-ene crosslinking reaction and establish a method for predicting thiol-ene reaction efficiency. Lastly, the feasibility of this hydrogel system for *in vitro* modeling was confirmed via assessment of metabolic activity of encapsulated primary human meniscal cells.

## 1. Introduction

Hydrogels are commonly used for biomedical applications, including modeling and tuning a diverse array of native extracellular matrix parameters for tissue engineering.^1,2^ Of specific interest is the ability to adjust hydrogel mechanical properties to understand how physical stimuli impact cell behavior independently of other microenvironmental signaling cues. Photoinitiated thiol-ene click chemistry is a well-documented method of synthesizing hydrogels with desired network properties to investigate mechanotransduction.^3^ This chemistry is popular due to the spatial and temporal control over the crosslinking reaction offered by photoinitiation, as well as benefits including rapid reaction rates, high yields, mild reaction conditions, and orthogonality.^3^ While most photoinitiated thiol-ene click chemistry reactions proceed with high fidelity and efficiency, there is also the potential for transition from a step-growth process to a chain-growth process,^3^ defined by homopolymerization of allyl groups rather than the reaction of allyl groups with thiols that defines step-growth reactions, or instances of unwanted side reactions such as the generation of disulfide bonds rather than thiol-ene crosslinks.^4^ A diverse array of polymeric backbones can be functionalized with a variety of different reactive -ene groups.^5–13^ Additionally, there are many options available for dithiol crosslinkers.^6,9,11,12^ The ability to select different polymers, types of reactive -ene groups, and crosslinking molecules makes photoinitiated thiol-ene crosslinked hydrogels highly customizable. The identity of these components, however, impact the thiol-ene reaction kinetics.^3,14^ Because of this, while photoinitiated thiol-ene click chemistry has historically been highly touted as efficient, recent studies have demonstrated that these reactions do not always proceed as predicted.

The use of hydrogels as tunable three-dimensional models of the cellular microenvironment requires a thorough understanding of crosslinked polymer network formation. The thiol-ene crosslinks formed during hydrogel synthesis not only determine the stiffness of the hydrogel system,^15,16^ but also determine hydrogel mesh size,^17^ defined as the average distance between neighboring crosslinked polymer chains in a swollen hydrogel.^17^ Mesh size determines the level of cell confinement, which can impact cell phenotype and function independently of hydrogel mechanics.^2,18^

We have previously developed a mechanically-tunable pentenoate-functionalized hyaluronic acid (PHA)-based hydrogel system for use as a two-dimensional *in vitro* model of knee meniscal tissue.^19^ This system is crosslinked via photoinitiated thiol-ene click chemistry and relies on modulating degree of functional group (i.e. reactive pentenoate -ene group) substitution (DoS) on the hyaluronic acid polymer backbone to tune crosslinking density and subsequently compressive storage modulus, or hydrogel stiffness. Using PHA hydrogels of increasing stiffness for two-dimensional culture of primary human meniscal cells, we demonstrated the ability to modulate meniscal cell morphology from spherical and balled up, which is potentially indicative of a chondrocyte-like phenotype, to spread with greater surface area, which is potentially indicative of a fibroblast-like phenotype. These results demonstrate a promising step towards the use of mechanical stimuli to control meniscal cell phenotype, a critical aspect of facilitating regeneration after meniscal damage. This original study only investigated the alteration of the degree of reactive -ene group substitution on hydrogel stiffness and did not evaluate the impact of other parameters (e.g. polymer concentration, crosslinking molar ratio, etc.) on PHA hydrogel network formation. Because of the unpredictable efficiency of photoinitiated thiol-ene click reaction, it is important to further characterize thiol-ene network formation for the PHA hydrogel system. This information is necessary for the use of the PHA hydrogel system for three-dimensional cell culture, where mesh size and cell confinement must be considered in addition to hydrogel mechanics, as well as to address the broader need to increase understanding of and identify pitfalls associated with thiol-ene crosslinked hydrogel network formation.

The objective of this study is to understand how crosslinking parameters including polymer concentration and crosslinker molar ratio (ratio of thiol groups to -ene groups) impact the photoinitiated formation of a thiol-ene crosslinked PHA hydrogel network and, subsequently, the network properties. This study will contribute to the long-term goal of expanding the applications of the PHA hydrogel system for three-dimensional *in vitro* modeling, as well as provide insight into the network formation of other thiol-ene crosslinked hydrogel systems. For this study, PHA DoS was held constant at ∼22% and polymer concentration and crosslinker thiol:-ene molar ratio were modulated (4, 6, or 8 PHA weight percent and 0.15:1, 0.5:1, or 1:1 molar ratio of thiol groups to reactive -ene groups) to demonstrate various methods of tuning hydrogel network properties (**Table 1**). The properties evaluated include plateau shear storage modulus measured via photorheology as well as shear storage modulus of hydrogels after crosslinking and swelling. The crosslinking reaction investigated in this study was simulated using Molecular Dynamics (MD) simulations to corroborate experimental results as well as develop a tool to predict the impact of compositional parameters on PHA hydrogel network formation. Relative hydrogel mesh size was also evaluated via fluorescence recovery after photobleaching (FRAP) to quantify diffusion coefficients. A relationship between storage modulus and relative mesh size for the PHA hydrogel system was established to demonstrate the potential to decouple hydrogel mesh size from hydrogel stiffness. Lastly, metabolic activity of encapsulated primary human meniscus cells was evaluated to confirm the feasibility of utilizing this PHA hydrogel system for three-dimensional cell culture and *in vitro* knee meniscus modeling.

**Table 1.** Summary of investigated hydrogel formulations.

|  |  | Crosslinker (PEG-DT) thiol:-ene molar ratio |  |  |
| --- | --- | --- | --- | --- |
|  |  | 1:1 | 0.5:1 | 0.15:1 |
| PHA concentration (wt% (w/v)) | 8 | 8 wt% 1:1 | 8 wt% 0.5:1 | 8 wt% 0.15:1 |
|  | 6 | 6 wt% 1:1 | 6 wt% 0.5:1 | 6 wt% 0.15:1 |
|  | 4 | 4 wt% 1:1 | 4 wt% 0.5:1 | 4 wt% 0.15:1 |

## 2. Results

### 2.1. Rheological characterization

#### 2.1.1. Photorheology

Photorheology results indicate that all investigated hydrogel formulations reach a steady state, evaluated by plateau shear storage modulus, within two minutes of UV light exposure at 365 nm and 6 mW/cm^2^ (**Figure 1 A-C**). Plateau shear storage modulus decreases predictably with decreasing polymer weight percentage when the molar ratio of crosslinker thiol groups to reactive -ene groups is held constant, indicating that decreasing polymer weight percentage results in decreased hydrogel stiffness (**Figure 1 D**). When polymer weight percentage is held constant, however, plateau shear storage modulus does not trend predictably with crosslinker molar ratio (**Figure 1 E**). Plateau shear storage modulus increases as crosslinker molar ratio increases from 0.15:1 to 0.5:1 thiol:-ene molar ratio and then decreases as crosslinker molar ratio increases from 0.5:1 to 1:1. This trend is seen across each of the weight percent groups investigated (8, 6, and 4 PHA weight percent). For reference, a crosslinker molar ratio of 1:1 indicates that every reactive -ene group present has one thiol group to react with after photoinitiated radical formation. For a thiol-ene click chemistry reaction with ideal efficiency, it would be expected that hydrogels fabricated with 1:1 molar crosslinking ratios would have a greater crosslinking density and therefore a greater storage modulus compared to hydrogels with 0.5:1 molar crosslinking ratios. Despite this, the formulations with 0.5:1 crosslinker molar ratios have increased plateau shear storage moduli compared to the formulations with 1:1 crosslinker molar ratios. Because interchain crosslinks (those in which a dithiol crosslinker molecule reacts with two -ene groups on two different PHA chains) are the thiol-ene connections that contribute to storage modulus, it can be deduced that the interchain crosslinking in the 0.5:1 molar ratio groups is likely proceeding more efficiently than that in the 1:1 molar ratio groups. Based on these experimental results, the crosslinking efficiency for this hydrogel system peaks somewhere between a molar crosslinking ratio of 0.15:1 thiol groups to -ene groups and 1:1 thiol groups to -ene groups.

**Figure 1.**
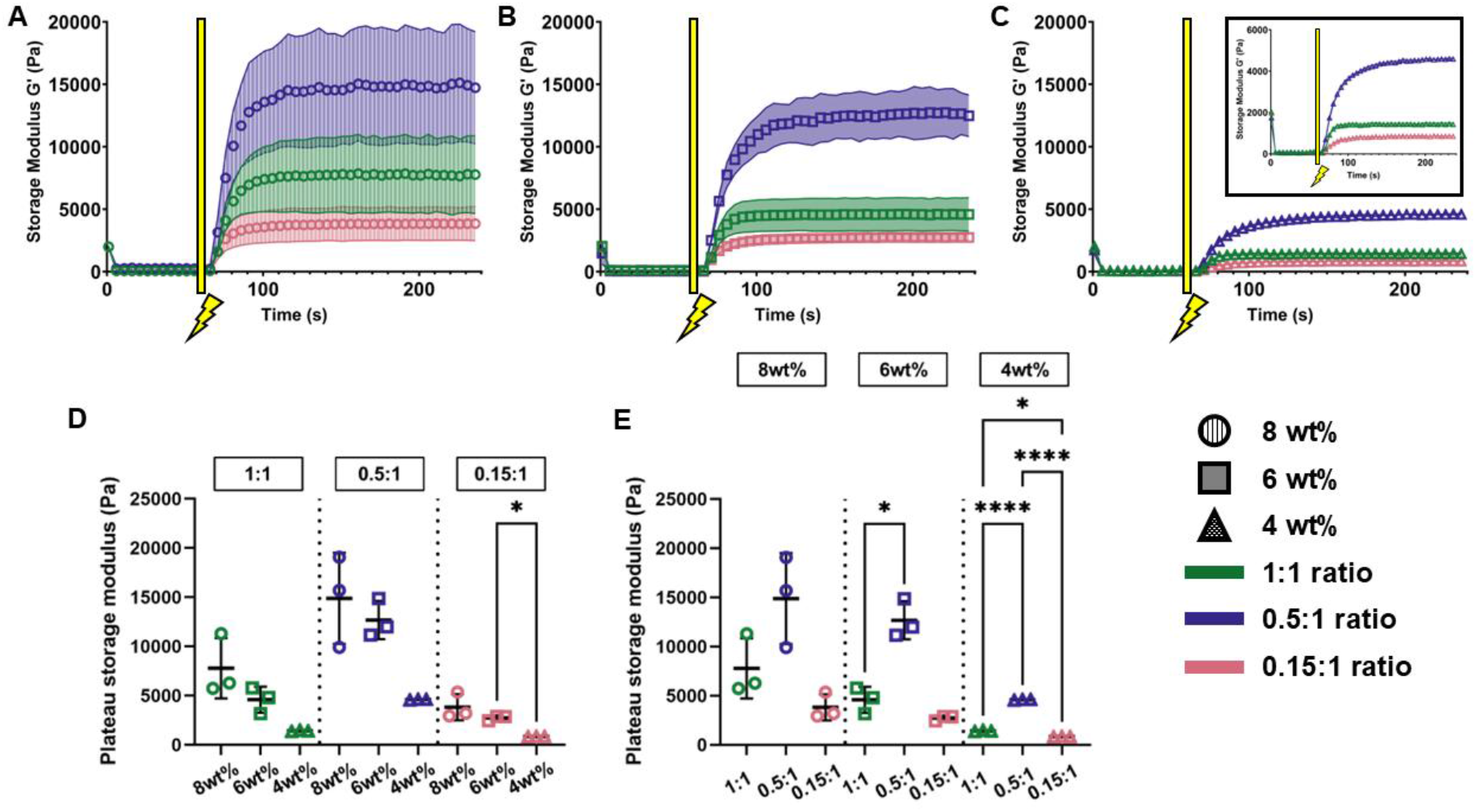
Photorheology results from A) 8 weight percent PHA formulations, B) 6 weight percent PHA formulations, and C) 4 weight percent PHA formulations (inset showing data on zoomed-in y-axis). Hydrogel precursor solution was conditioned for 60 seconds before UV light was turned on (signified by lightning bolt icon). Average plateau storage moduli for formulations grouped by thiol:-ene molar ratio (D) and by polymer concentration (E), determined by taking an average of storage moduli values from 200 s to 240 s. In panels A-C, icons represent the average of 3 runs and the shaded portion represents the standard deviation. In panels D-E, * indicates p≤0.05, ** indicates p≤0.01, *** indicates p≤0.001, and **** indicates p≤0.0001.

#### 2.1.2. Post-swelling strain amplitude sweeps

While photorheological characterization is an important tool for the evaluation of photoinitiated crosslinking kinetics, it is necessary to investigate mechanical properties of this PHA hydrogel system after crosslinking is complete and hydrogels have undergone swelling equilibration. Equilibrated hydrogels will be used for *in vitro* experiments, so post-swelling rheological characterization data is necessary for application of this PHA hydrogel system to cell studies.

Rheological characterization after the equilibration of hydrogel samples via swelling in PBS (**Figure 2**) also demonstrates that shear storage modulus decreases with decreasing polymer weight percentage when the molar ratio of crosslinker thiol groups to reactive -ene groups is held constant (**Figure 2 A-C and D**), confirming that decreasing polymer weight percentage results in decreased hydrogel stiffness. Interestingly, the relationship between crosslinker molar ratio and shear storage modulus when polymer weight percent is held constant, however, does not trend the same way in equilibrated hydrogel samples compared to pre-equilibrated hydrogel samples (i.e. those characterized with photorheology) (**Figure 2 E**). For 8 weight percent groups of equilibrated hydrogels, average shear storage modulus increases as crosslinker molar ratio increases from 0.15:1 to 0.5:1 and then decreases as crosslinker molar ratio increases from 0.5:1 to 1:1. This is the same trend seen across all investigated polymer weight percentage groups when characterized with photorheology. For the 6 weight percent groups of equilibrated hydrogels, average shear storage modulus increases as crosslinker molar ratio increases from 0.15:1 to 0.5:1 and then exhibits no difference as crosslinker molar ratio increases from 0.5:1 to 1:1. Lastly, for the 4 weight percent groups of equilibrated hydrogels, average shear storage modulus increases as crosslinker molar ratio increases from 0.15:1 to 1:1.

**Figure 2.**
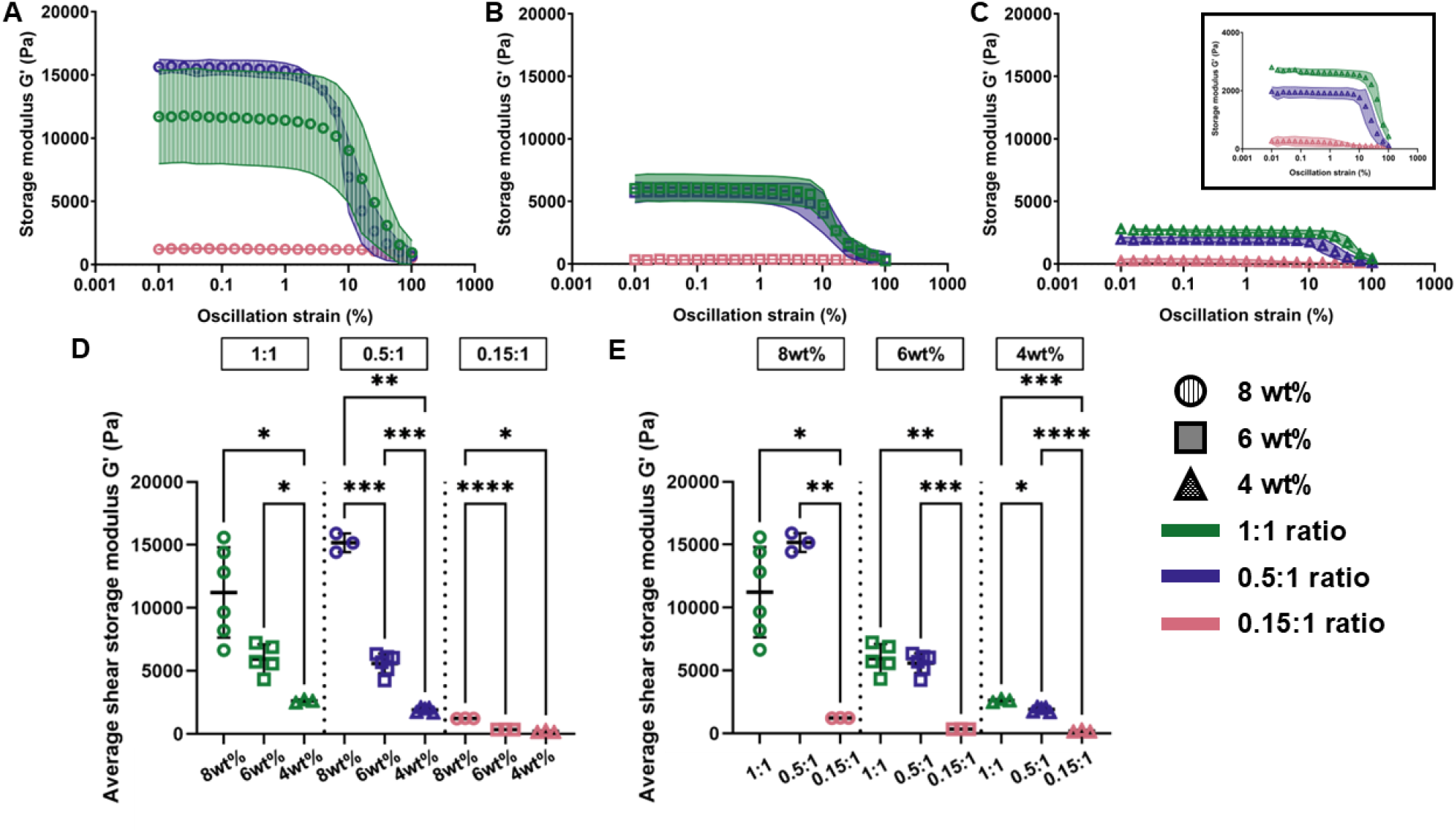
Post-swelling rheology results from A) 8 weight percent PHA formulations, B) 6 weight percent PHA formulations, and C) 4 weight percent PHA formulations (inset showing data on zoomed-in x-axis). Strain amplitude sweeps were conducted from 0.01% to 100% strain. Average shear storage moduli for formulations grouped by thiol:-ene molar ratio (D) and by polymer concentration (E), determined by taking an average of shear storage moduli values from 0.01% strain until onset point strain value. In panels A-C, icons represent the average of 3 runs and the shaded portion represents the standard deviation. In panels D-E, * indicates p≤0.05, ** indicates p≤0.01, *** indicates p≤0.001, and **** indicates p≤0.0001.

When directly comparing the pre-equilibration plateau shear storage modulus values for each PHA hydrogel formulation to the average shear storage modulus values after equilibration for each corresponding formulation (**Figure 3**), differences in swelling behavior that contribute to resistance to deformation can be seen. Equilibration of PHA hydrogel samples in PBS after photoinitiated crosslinking causes increases in average shear storage moduli for all formulations with a 1:1 crosslinker molar ratio. On the other hand, equilibration in PBS causes decreases in average shear storage moduli for all hydrogels with formulations with a 0.15:1 crosslinker molar ratio. Equilibration in PBS also causes decreases in average shear storage moduli of hydrogels with the 6 weight percent, 0.5:1 crosslinker molar ratio formulation and with the 4 weight percent, 0.5:1 crosslinker molar ratio formulation. For hydrogels with the 8 weight percent, 0.5:1 crosslinker molar ratio formulation, swelling of the hydrogel samples in PBS does not significantly change the average shear storage modulus. These differences may be due to a variety of factors. Increasing the concentration of the very hydrophilic PHA polymer may increase degree of swelling,^20,21^ though uptake of water into the hydrogel may not contribute to deformation resistance to the same degree that interchain crosslinks formed in each hydrogel contribute to deformation resistance. Additionally, photorheology experiments indicate that the PHA hydrogel crosslinking reaction is not 100% efficient, indicating that some material remains unreacted but constrained within the hydrogel during photorheological characterization. This unreacted material is likely washed out of the hydrogel during equilibration, changing the hydrogel’s ability to resist deformation. On the other hand, in this hydrogel system, the length of the PHA polymers far exceeds the crosslinker length, indicating that these PHA polymers are more likely to have large decreases in polymer chain entanglement immediately after network formation, especially at higher concentrations of PHA. This could also explain the relationships shown in **Figure 3**.

**Figure 3.**
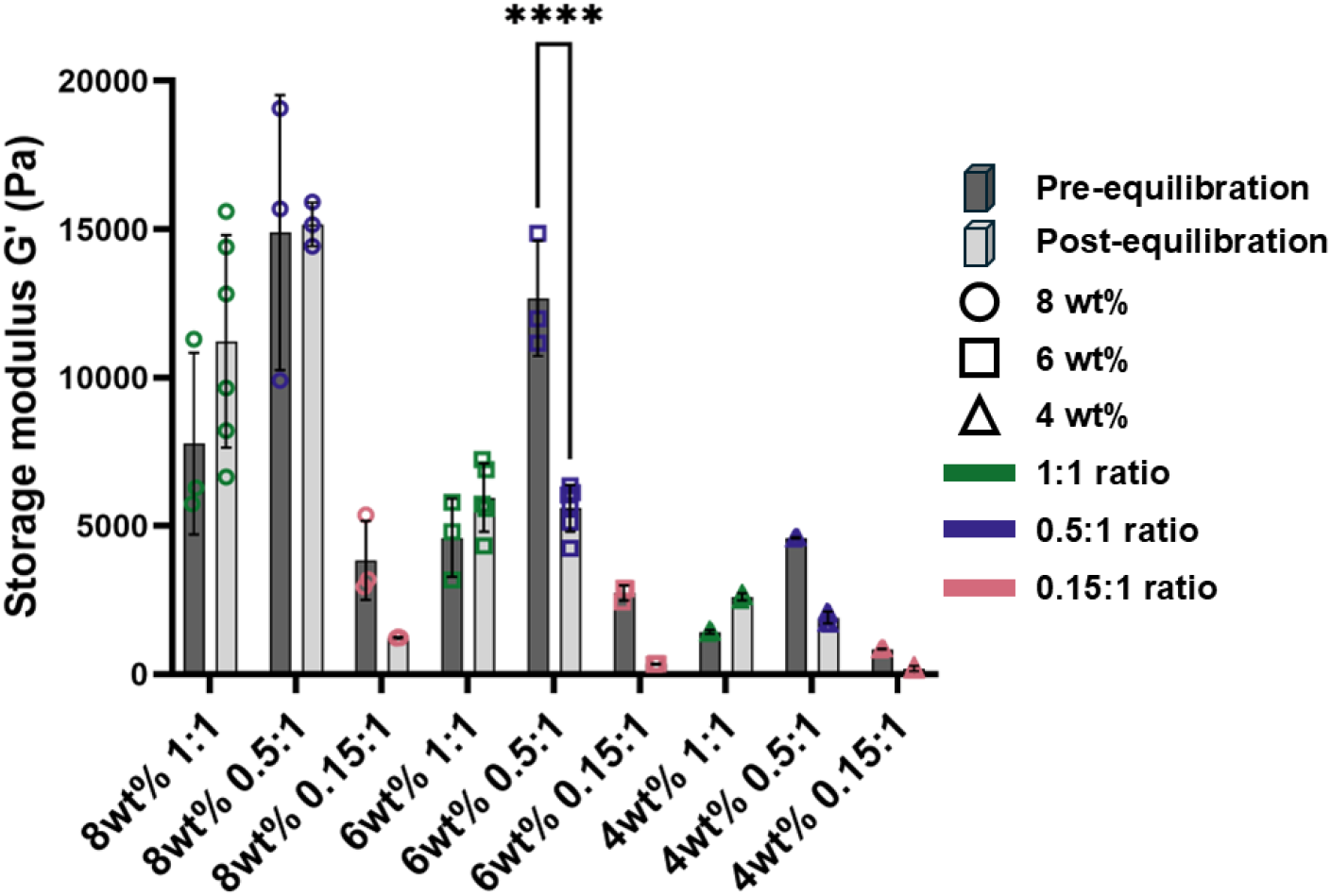
Comparison of pre-equilibration shear storage moduli (determined via photorheology) and post-equilibration shear storage moduli (determined via post-swelling strain amplitude sweeps). Pre-equilibration values shown in dark grey and pre-equilibration values shown in light grey. For pre-vs. post-equilibrium comparisons, * indicates p≤0.05, ** indicates p≤0.01, *** indicates p≤0.001, and **** indicates p≤0.0001.

### 2.2. Molecular Dynamics simulations

The loss of thiol-ene crosslinking reaction efficiency demonstrated here as well as in other studies^7,8,12,22^ can be attributed to a lack of formation of interchain crosslinks relative to the number of available thiol groups and available -ene groups. It is posited that other undesired reactions that do not contribute to the formed hydrogel’s resistance to deformation are also present. These include intrachain crosslinks in which a dithiol crosslinker molecule bonds to two -ene groups on the same polymer chain forming a loop, disulfide bridge formation in which two thiol groups form a bond,^4^ and pendant thiol formation in which only one thiol group on a dithiol crosslinking molecule reacts with an -ene group leaving the other thiol group unreacted.^23^ Additionally, there is a possibility that some -ene groups and dithiol crosslinker molecules are left entirely unreacted. To evaluate the types of thiol-ene reactions occurring in the PHA hydrogel system presented here, as well as to predict the thiol:-ene ratio at which maximum crosslinking efficiency is achieved, the present thiol-ene reaction was simulated using Molecular Dynamics (MD) simulations.

All carbon-sulfur bonds formed in thiol-ene reactions were tracked and organized using MD simulations. For these simulations, total thiol reaction efficiency was defined as the ratio of the number of reacted thiol groups to the total number of thiol groups available in the system. The thiol reaction efficiency for each type of reaction (interchain crosslink, intrachain crosslink, and pendant or incomplete reaction, shown in **Figure 4 A**) is defined as the ratio of the number of reacted thiol groups present in each type of reaction to the total number of thiol groups available in the system. The simulated reaction efficiencies for the presented hydrogel system at thiol:-ene molar ratios of 0.5:1, 0.77:1, and 1:1 are summarized in **Figure 4 B**. The total thiol reaction efficiency as well as the efficiencies for interchain crosslink formation, intrachain crosslink formation, and pendant reaction for each thiol:-ene molar ratio are presented as a function of MD simulation time in **Figure 4 C-F**. Notably, the total thiol reaction efficiency (**Figure 4 C**) is maximized at a thiol:-ene molar ratio of 0.5:1, meaning that thiol-ene reactions of any kind (interchain, intrachain, or pendant) occur at a greater efficiency in hydrogel systems with available thiol groups totaling half the number of available -ene groups than in hydrogel systems more available thiol groups relative to available -ene groups. The next most efficient ratio simulated is 0.77:1 and the least efficient ratio simulated is 1:1. When looking at interchain crosslink reaction efficiency only (**Figure 4 D**), the same trend is true: efficiency is maximized in the 0.5:1 ratio hydrogel crosslinking simulations and the 1:1 ratio results in the lowest efficiency. Because interchain crosslinking contributes most to resultant hydrogel mechanical properties including compressive or shear storage modulus, these MD simulations confirm our experimental results and help to narrow the range of possible thiol:-ene ratios at which maximum interchain crosslinking is achieved for this system to 0.15:1 to 0.77:1. When looking at intrachain crosslink reaction efficiency (or loop formation) only (**Figure 4 E**), the same trend continues to hold true but the differences in efficiency between the simulated ratios are much smaller. Lastly, when looking at pendant formation efficiency only (**Figure 4 F**), the same trend continues to hold true except that the efficiencies for the 0.77:1 and 1:1 ratios are largely similar. Of all the possible types of thiol reactions that can form during thiol-ene crosslinking in this simulated hydrogel system, the smallest share of formed thiol-ene bonds fall into the category of intrachain crosslinks. Interestingly, for the 0.5:1 and 0.77:1 thiol:-ene molar ratio groups, the share of formed thiol-ene bonds in the interchain crosslink category is roughly equal to the share in the pendant formation category. The hydrogel system with a ratio of 1:1, however, forms pendant thiols with greater efficiency than it forms interchain crosslinks, further illustrating the inefficiency of the system at a thiol:-ene molar ratio of 1:1 as assessed in this simulation field. If the intent of the mechanically tunable hydrogel system is to maximize mechanical properties including compressive or shear storage modulus, these simulations can be used to determine the thiol:-ene ratio that provides maximum efficiency of interchain crosslink formation. It may, however, be desirable to modulate hydrogel mechanical properties while also controlling the amount of pendant thiol present after photoinitiated crosslinking is complete for downstream modification of the hydrogel network.^6,23^ In that case, this simulation is also useful to help predict the efficiency of pendant thiol group formation relative to the efficiency of interchain crosslinking.

**Figure 4.**
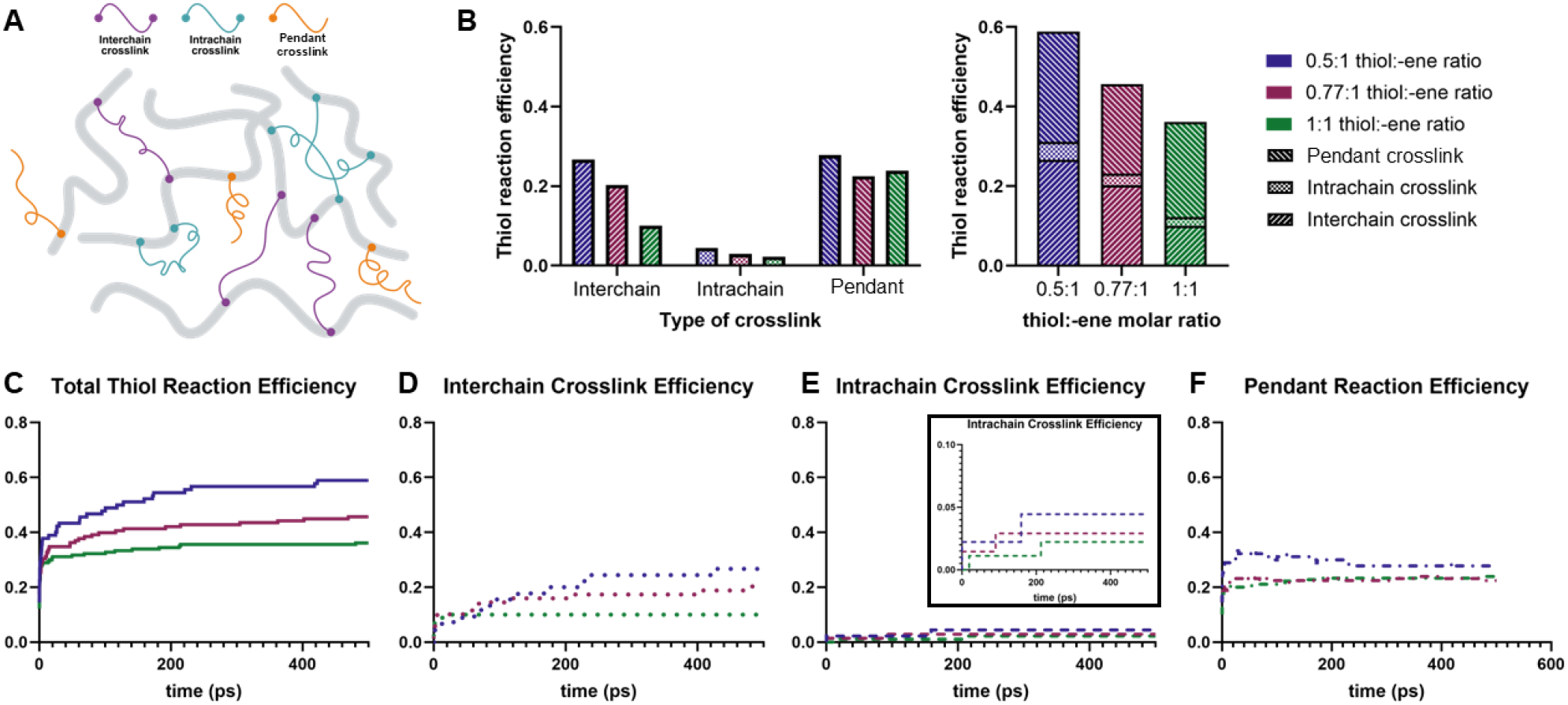
Molecular Dynamics simulations results. A) Schematic showing types of reactions simulated including interchain, intrachain, and pendant (incomplete) crosslinks, B) Thiol reaction efficiency, defined as the ratio of the number of reacted thiol groups to the total number of thiol groups available for reaction, separated by type of reaction for each simulated thiol:-ene molar ratio, C-F) Thiol reaction efficiency over time from the beginning of the reaction for total thiol reactions (C), interchain crosslinks (D), intrachain crosslinks (E), and pendant (incomplete) crosslinks (F). Data analysis performed by Yiwen Zheng.

Of note, the maximum total thiol reaction efficiency for this simulation is less than 60%, meaning that more than 40% of the thiol groups present during PHA hydrogel crosslinking remain unreacted. This may be attributed to the use of polymer concentrations below the critical overlap concentration (c*) for hyaluronic acid. The critical overlap concentration (c*) is the concentration at which polymer chains in a solution begin to interact or entangle. At polymer concentrations below c*, polymer chains are separated and do not interact, which may decrease the efficiency of crosslinking, or the total thiol reaction efficiency. For hyaluronic acid with a molecular weight of 60 kDa, the critical overlap concentration is greater than 8,600 μg/cm^3^ which is equivalent to approximately 8.6 weight percent.^24^ Because this study involved PHA concentrations below the critical overlap concentration, the lack of polymer chain interaction may explain the unexpected efficiency results.

### 2.3. Fluorescence recovery after photobleaching

Fluorescent recovery after photobleaching (FRAP) was used to calculate the diffusion coefficient of green fluorescent protein (GFP) throughout equilibrated PHA hydrogels synthesized using each of the 9 investigated formulations. **Figure 5 A** shows representative pre-bleach, bleach, and recovery images for PHA hydrogel formulations with the highest and lowest stiffness based on photorheology results (**Figure 1**) and post-swelling rheological characterization (**Figure 2**). The calculated diffusion coefficients are also shown in **Figure 5**. When the molar ratio of crosslinker thiol groups to reactive -ene groups is held constant, diffusion coefficient of GFP increases with decreasing polymer weight percent (**Figure 5 B**). An increased diffusion coefficient indicates an increased rate of diffusion for GFP through the hydrogel, which indicates a larger hydrogel mesh size. A larger hydrogel mesh size indicates a decreased density of crosslinks. A decrease in crosslink density typically indicates a decreased in mechanical properties including shear storage moduli and compressive storage moduli. Therefore, an increase in diffusion coefficient is associated with a decrease in storage moduli. This data is aligned with rheological characterization results for PHA hydrogels both before and after equilibration of PHA hydrogels via swelling (**Figure 1 and 2**): when thiol:-ene molar ratio is held constant and polymer weight percent decreases, GFP diffusion coefficient increases and storage modulus decreases.

**Figure 5.**
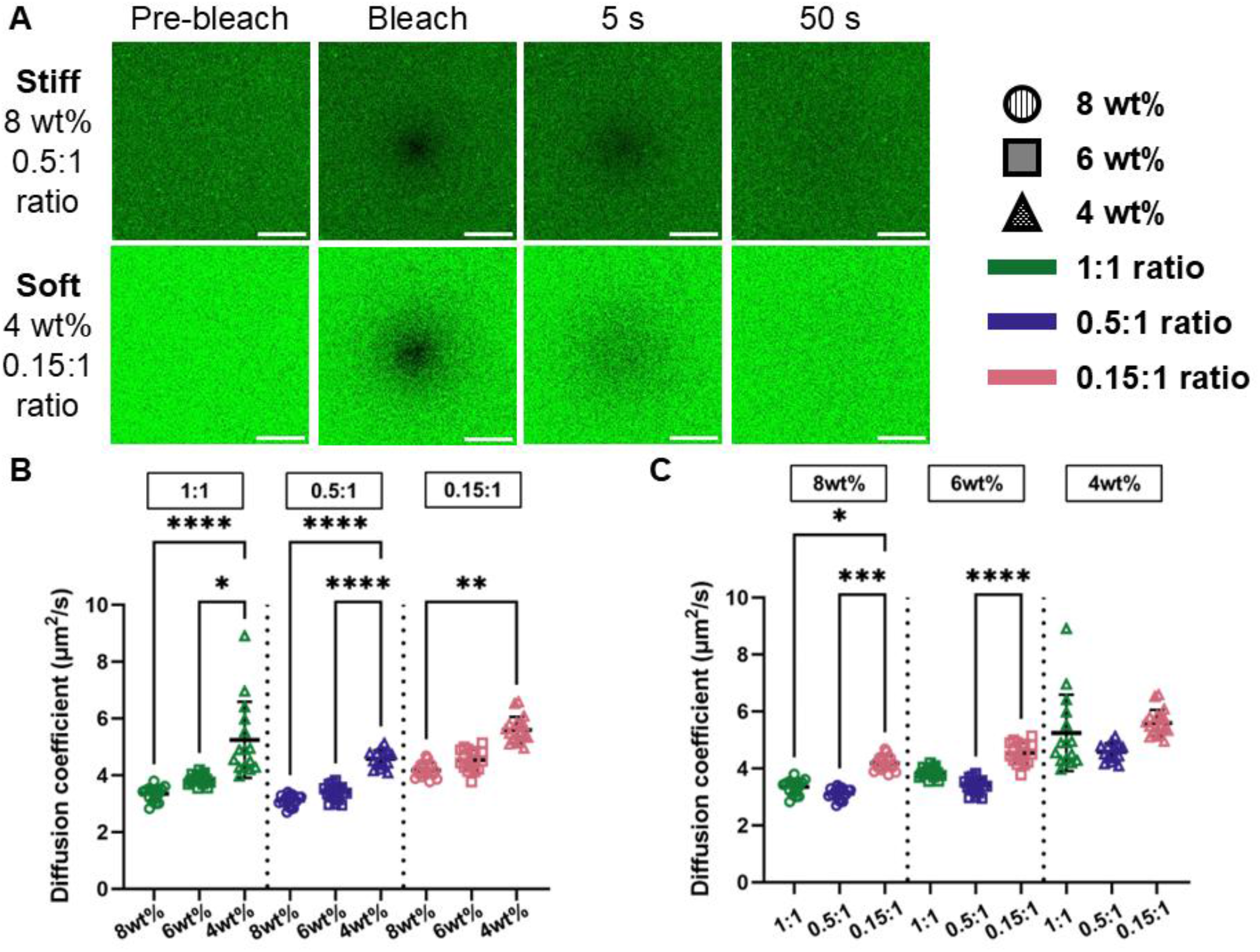
FRAP results. A) Representative FRAP images showing the bleach spot and the recovery of fluorescence intensity after bleaching as a function of time, B-C) Diffusion coefficient of GFP through the PHA hydrogel system for all 9 investigated formulations grouped by either thiol:-ene molar ratio (B) or polymer weight percent (C). In panels B-C, * indicates p≤0.05, ** indicates p≤0.01, *** indicates p≤0.001, and **** indicates p≤0.0001.

When polymer weight percent is held constant and thiol:-ene molar ratio is modulated, however, the GFP diffusion coefficient trend matches only the pre-equilibration shear storage moduli data (**Figure 1**) and not the post-equilibrium shear storage moduli data. Specifically, as thiol:-ene molar ratio increases from 0.15:1 to 0.5:1, the GFP diffusion coefficient calculated via FRAP decreases (**Figure 5 C**), predicting increased mechanical properties based on the relationship between diffusion, hydrogel mesh size, hydrogel crosslinking density, and resulting mechanical properties. This predicted increase in mechanical properties is confirmed when looking at both pre-equilibration photorheology data (**Figure 1**) and post-equilibration rheology data (**Figure 2**). The difference is seen as thiol:-ene molar ratio increases from 0.5:1 to 1:1. This increase in molar ratio results in increased diffusion coefficients for all investigated polymer weight percents, predicting decreased mechanical properties. This predicted decrease in mechanical properties is seen across all investigated polymer weight percents when shear storage modulus was calculated prior to swelling using photorheology (**Figure 1**), but it is only seen in the 8 weight percent groups when shear storage modulus was calculated after equilibration via swelling using post-swelling strain amplitude sweeps (**Figure 2**). Instead, in the 6 weight percent groups, post-swelling shear storage modulus remains the same as thiol:-ene molar ratio is increased from 0.5:1 to 1:1, and in the 4 weight percent groups, post-swelling shear storage modulus increases as thiol:-ene molar ratio is increased from 0.5:1 to 1:1 (**Figure 2 E**). This is interesting because FRAP data is collected on equilibrated PHA hydrogels and it would be expected that an increase in GFP diffusion coefficient would be associated with a decrease in post-swelling shear storage modulus.

When plotting GFP diffusion coefficient as a function of pre-equilibration shear storage modulus, or plateau storage modulus (**Figure 6 A**), and, separately, as a function of post-equilibration shear storage moduli (**Figure 6 B**), it is clear that the hypothesized relationship between hydrogel mesh size and crosslinking density does not necessarily hold true across all investigated formulations of the PHA hydrogel system.

**Figure 6.**
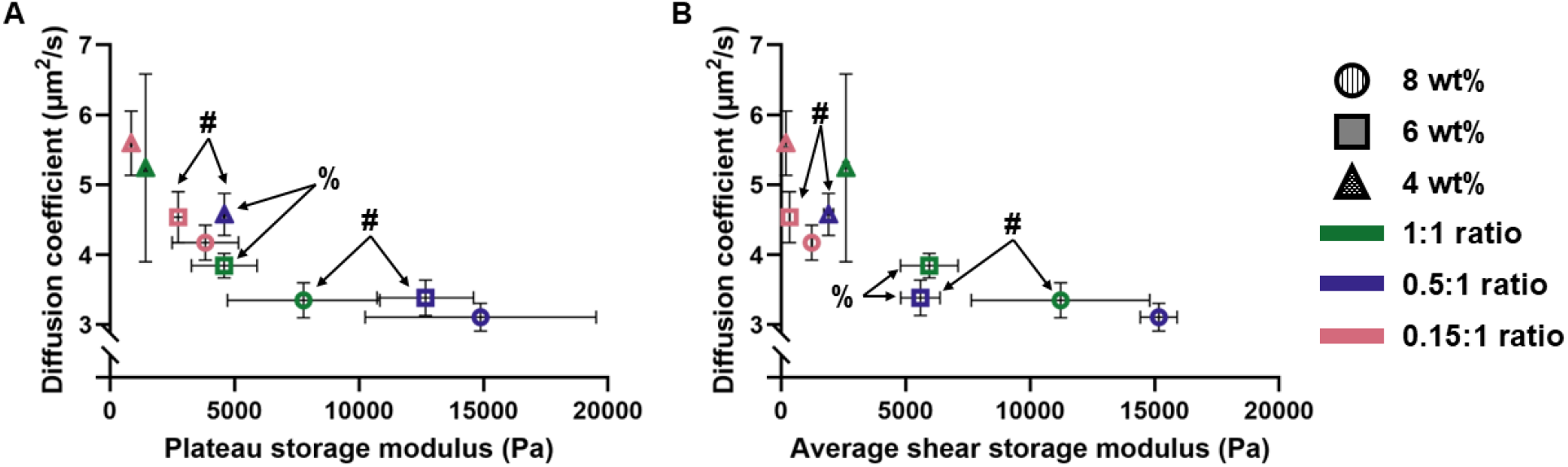
GFP diffusion coefficient determined via FRAP analysis as a function of either A) pre-equilibration hydrogel shear storage modulus or B) post-equilibration hydrogel shear storage modulus for all 9 PHA hydrogel formulations investigated. **%** denotes formulations in which stiffness is similar but diffusion coefficient, and subsequently mesh size, is different, and **#** denotes formulations in which diffusion coefficient, and subsequently mesh size, is similar but stiffness is different.

Interestingly, plotting GFP diffusion coefficient as a function of hydrogel shear storage modulus demonstrated a potential opportunity to decouple hydrogel mesh size and hydrogel mechanics. As discussed above, hydrogel mesh size is indicative of hydrogel crosslinking density which is typically inextricably linked to hydrogel mechanics. This makes it incredibly difficult to investigate the impact of microenvironment stiffness separately from the impact of physical confinement on cell behavior when cells are encapsulated in hydrogels for three-dimensional cell culture. Within the nine formulations of PHA hydrogels investigated, there are two examples of this decoupling in equilibrated hydrogels, shown in **Figure 6 B**. PHA hydrogels with the formulations 6 weight percent, 0.5:1 thiol:-ene molar ratio and 6 weight percent, 1:1 thiol:-ene molar ratio appear to have similar shear storage moduli, but different GFP diffusion coefficients and therefore network mesh sizes. These formulations represent an opportunity to investigate the impacts of differential levels of cell confinement on encapsulated cells while maintaining a constant substrate mechanics. On the other hand, PHA hydrogels with the formulations 6 weight percent, 0.5:1 thiol:-ene molar ratio and 8 weight percent, 1:1 thiol:-ene molar ratio appear to have similar GFP diffusion coefficients and therefore network mesh sizes, but different shear storage moduli. These formulations represent an opportunity to investigate the impacts of substrate mechanics on encapsulated cells while maintaining a constant level of cell confinement.

### 2.3. Cell encapsulation and metabolic activity

Primary human meniscal cells were encapsulated in PHA hydrogels with two different formulations (6 weight percent, 0.5:1 thiol:-ene molar ratio and 8 weight percent, 1:1 thiol:-ene molar ratio) to investigate metabolic activity via PrestoBlue assay (**Figure 7**). Over the course of 5 days, the metabolic activity of primary human meniscal cells encapsulated in PHA hydrogels increased and then held at a roughly constant level from day 5 to day 7. This provides evidence for the viability of encapsulated cells cultured in the PHA hydrogel system and therefore the feasibility of the PHA hydrogel system as a mechanically tunable *in vitro* model to investigate the effects of microenvironment mechanics and cellular confinement by hydrogel mesh size on cell behavior. At all timepoints except for day 3, the metabolic activity was similar between the two PHA hydrogel formulations investigated. At day 3, however, the metabolic activity of cells encapsulated in the hydrogels with the 8 weight percent, 1:1 thiol:-ene molar ratio formulation was greater than the metabolic activity of cells encapsulated in the hydrogels with the 6 weight percent, 0.5:1 thiol:-ene molar ratio formulations. As discussed above, these hydrogel formulations have similar mesh size (i.e. similar GFP diffusion coefficients) but the 8 weight percent, 1:1 thiol:-ene molar ratio formulation had a greater average shear storage modulus compared to the 6 weight percent, 0.5:1 thiol:-ene molar ratio formulation, indicating it is a stiffer hydrogel formulation. This difference in stiffness may be the reason for the demonstrated differences in metabolic activity. While further studies are needed to confirm this hypothesis, FRAP analysis of diffusion coefficient helps to exclude differential levels of cell confinement as a reason for the day 3 metabolic activity differences.

**Figure 7.**
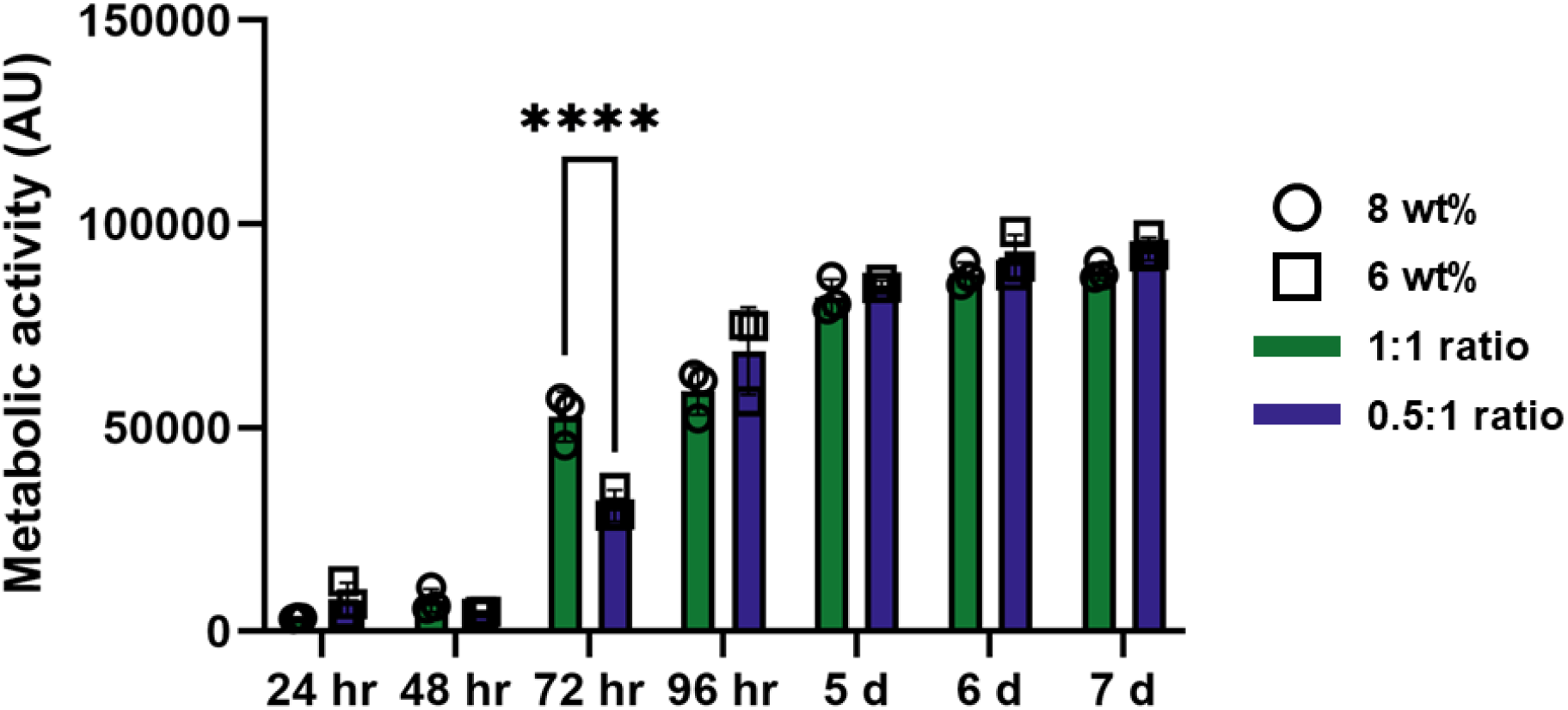
Metabolic activity of primary human meniscal cells encapsulated in PHA hydrogels and cultured in 3D over the course of 1 week. For formulation comparisons, * indicates p≤0.05, ** indicates p≤0.01, *** indicates p≤0.001, and **** indicates p≤0.0001.

## 3. Discussion

The work presented here investigated the impact of both polymer concentration and crosslinker molar ratio on PHA thiol-ene hydrogel network formation and resulting hydrogel mechanics and hydrogel mesh size. The most predictable trend appreciated from this work is the one characterizing the relationship between polymer weight percent and hydrogel mechanics and mesh size. When thiol:-ene molar ratio is held constant, an increase in polymer weight percent results in hydrogels with increased mechanics in terms of both pre- and post-equilibration shear storage modulus as well as decreased mesh size and therefore decreased GFP diffusion coefficient. This trend is consistent with that seen in other hyaluronic acid-based systems that utilize thiol-ene crosslinking chemistry,^7^ as well as poly(ethylene glycol)-based systems that utilize thiol-ene crosslinking chemistry.^6^

The relationship between thiol:-ene molar ratio and hydrogel mechanics and mesh size, however, is less predictable, especially when comparing hydrogels before and after equilibration via swelling. The idea that the number of thiol:-ene crosslinks formed during hydrogel synthesis peaks somewhere between thiol:-ene molar ratios of 0.15:1 and 1:1 rather than increasing linearly as thiol:-ene molar ratio increases is not necessarily novel. This phenomenon has been documented in other hydrogel studies using thiol-ene click chemistry for crosslinking, but peak efficiency is dependent on reaction details including polymeric backbone identity and concentration. Norbornene functional groups are one of the most common functional groups for the thiol-ene crosslinking reaction; the relationship between thiol:-ene molar ratio and crosslinking reaction efficiency has not been documented for hydrogel systems involving pentanoate functional groups. For norbornene-functionalized hyaluronic acid hydrogels crosslinked with dithiothreitol, hydrogel storage modulus has been maximized at a ratio of 0.6:1 thiol groups to -ene groups,^7^ and it has been shown in this system that increasing the ratio beyond 1:1 results in decreasing hydrogel storage modulus, indicating that an excess of thiol groups may negatively impact thiol-ene reaction efficiency. For hydrogels composed of carbocymethyl cellulose modified with norbornene groups and crosslinked with dithiothreitol, however, hydrogel compressive modulus increases predictably with increasing molar ratios of thiol groups to -ene groups from 0.25:1 to 1:1.^12^ Upon the inclusion of cellulose nanofibrils, it was determined that a thiol:-ene molar ratio of less than 1:1 maximized hydrogel compressive modulus. For hydrogels involving a crosslinking reaction between norbornene-functionalized poly(ethylene glycol) and poly(ethylene glycol)-dithiol, compressive modulus increases predictably with increasing thiol:-ene molar ratios between 0.25:1 and 1:1.^6^ These studies also vary in other crosslinking parameters, including polymer weight percent, photoinitiator identity, crosslinker identity, and irradiation time, further illustrating that thiol-ene reaction efficiency is system-dependent and must be characterized for each individual system.

The MD simulations performed in this work may provide a more efficient method of characterizing thiol-ene reaction efficiency for individual hydrogel systems. The MD simulations shown here map to the collected experimental data as well as provide insight into the crosslinking efficiency of PHA hydrogel formulations that were not part of the experimental investigation. Importantly, while MD simulations confirm experimental data in identifying that a thiol:-ene ratio of 0.5:1 results in peak crosslinking efficiency, this crosslinking efficiency is still relatively low at only about 60%, meaning that of all available thiol groups in the system, only 60% react with an - ene group. While some may form disulfide bonds, a possibility that was not accounted for in these MD simulations, this efficiency is still relatively low given the highly touted efficiency of thiol-ene click chemistry in general. From these results, one can attribute this to the use of polymer concentrations below the critical overlap concentration of 60 kDa hyaluronic acid or deduce that it is likely that unreacted molecules (e.g. crosslinker or photoinitiator) remain in the PHA hydrogel after the crosslinking reaction equilibrates. These unreacted molecules likely wash out of the hydrogel during swelling or equilibration.

The presence of unreacted molecules in the PHA hydrogel before equilibration and the lack of these unreacted molecules after equilibration may provide some explanation for the differences in hydrogel mechanics seen before and after hydrogel swelling if the unreacted molecules are contributing to the deformation of shear force. This may be the case for all 0.15:1 thiol:-ene molar ratio formulations, where equilibration of PHA hydrogels in PBS decreased shear storage modulus and therefore hydrogel stiffness, as well as for the 6 weight percent, 0.5:1 thiol:-ene molar ratio group and the 4 weight percent, 0.5:1 thiol:-ene molar ratio group, where swelling also caused a decrease in hydrogel shear storage modulus. This logic doesn’t apply well to the effects of swelling on all 1:1 thiol:-ene molar ratio formulations, where equilibration of the PHA hydrogels in PBS increased shear storage modulus and therefore hydrogel stiffness. This is curious because Molecular Dynamics simulations indicate a decreased crosslinking efficiency in the 1:1 thiol:-ene molar ratio formulations relative to 0.5:1 thiol:-ene molar ratio formulations, indicating that there should be an increased presence of unreacted molecules in 1:1 thiol:-ene molar ratio hydrogels compared to 0.5:1 thiol:-ene molar ratio hydrogels. Overall, these results demonstrate the unpredictable nature of swelling in PHA hydrogels, and likely other hydrogel systems, with different crosslinking molar ratios and different degrees of crosslinking. Because hydrogels must be equilibrated for use as *in vitro* modeling, it is important to evaluate hydrogel mechanics after equilibration in solutions of interest (e.g. PBS or cell culture media) rather than relying on pre-swelling shear storage moduli such as that collected using photorheology.

The work presented here also shows a potential decoupling of hydrogel mesh size from hydrogel mechanics, two hydrogel network properties that have historically been closely tied together. Many studies have shown simultaneous modulation of hydrogel mechanics and hydrogel mesh size,^25,26^ but only a few have shown promise of independent tuning of these properties. One such study incorporated secondary reactive species into poly(ethylene glycol) (PEG) diacrylate (PEGDA) hydrogels to obtain hydrogels with varying moduli and consistent average mesh size.^27^ Another study, also using PEG-based hydrogels with acrylate crosslinking, incorporated small amounts of four-arm PEGDA crosslinker to increase hydrogel moduli without significantly impacting hydrogel mesh size.^28^ Neither of these studies, however, investigated the impact of hydrogel moduli on cell behavior independently of the impact of hydrogel mesh size (and vice versa). A third study, however, did independently investigate the impact of these parameters on encapsulated cells. In this case, another PEG-based hydrogel was utilized and the hydrogel modulus and mesh size were decoupled by tuning both the molecular weight and the concentration of the polymer.^29^ These parameters independently modulated the viability, proliferation, and differentiation of oligodendrocyte precursor cells when encapsulated in the hydrogels.^29^ The study presented here only looks at metabolic activity of primary human meniscal cells encapsulated PHA hydrogels of different stiffnesses but similar mesh size, but one can still appreciate that the metabolic activity of these cells can be altered by stiffness of their microenvironment independent of their level of confinement in the 3D crosslinked polymer network.

## 4. Limitations

Although the work presented in this study is robust, several limitations impact the conclusions drawn and the future directions. MD simulations were performed using a condensed model of the PHA hydrogel crosslinking reaction. Specifically, the molecular weights of the PHA polymers and the PEG-DT crosslinking molecules were decreased as well as the number of PHA polymers and PEG-DT crosslinking molecules available in the system. This was done to simplify the simulation and ensure that a sufficient amount of crosslinks were formed within a reasonable simulation timeframe. It is recognized that this may skew the resultant crosslinking efficiency data shown in this study. Additionally, the MD simulations were only able to account for carbon-sulfur bonds and not sulfur-sulfur bonds, so the presence of disulfide bond formation could not be accounted for using these simulations.

Additionally, FRAP is a useful but limited method of evaluating hydrogel mesh size because it can only approximate relative diffusion of GFP through a hydrogel system; this technique is unable to provide any quantitative measurement of hydrogel mesh size. Despite this, FRAP analysis is still useful for understanding the relationship between hydrogel mesh size and hydrogel mechanics, including the potential to decouple these parameters for independent investigation.

## 5. Conclusions

Overall this study provides increased insight into the network formation of the PHA hydrogel system, providing extensional knowledge of other thiol-ene click-chemistry based hydrogel systems in general. This work demonstrates the ability to tune hydrogel mechanics and mesh size using both polymer concentration and thiol:-ene crosslinking molar ratio, as well as establishes a method of simulating thiol:-ene crosslinking in the PHA hydrogel system using MD simulations to predict resultant mechanical properties without the need for experimentation. This study also highlights the importance of hydrogel equilibration and the complex relationships between crosslinking density, crosslinking efficiency, and polymer hydrophilicity as well as the impact of these relationships on hydrogel mechanics before and after equilibration. Lastly this work demonstrates that the PHA hydrogel system is a feasible system for three-dimensional culture of encapsulated cells and *in vitro* modeling of the cellular extracellular matrix, as well as unveils a potential method of decoupling hydrogel mechanics from hydrogel mesh size for the independent investigation of the impact of these parameters on cells cultured in three-dimensions.

## 6. Materials and Methods

### 6.1. Functionalization and preparation of PHA

PHA was synthesized via an esterification reaction as previously described^19^ (**Figure 8**). All steps of the functionalization reaction were performed at 4°C. Briefly, sodium hyaluronate (Lifecore Biomedical, MW ∼60 kDa) was dissolved overnight in DI water at 20 mg/mL. The sodium hyaluronate solution was then transferred to a tri-neck round-bottom flask and combined with N,N-dimethylformamide (DMF) (Millipore Sigma) at a volumetric ratio of 3:2 sodium hyaluronate solution:DMF. The solution pH was adjusted to 9.0 ± 0.1 by addition of 1 M NaOH (Millipore Sigma) and monitored by pH probe throughout the reaction. 4-pentenoic anhydride (PA) was added to the reaction in 50-µL increments to achieve a molar ratio of 1:1 sodium hyaluronate:PA. Using 1 M NaOH, the reaction pH was maintained within the range of 8.0 to 9.0 for ∼5 hours, starting after all PA was added. After ∼5 hours of monitoring, the reaction pH was adjusted to 9.5 and left to stir for an additional ∼19 hours (24 hours total). After the completion of the reaction, the product was dialyzed using 6-8 kDa MWCO tubing (Spectrum) and DI water for at least 4 hours. The DI water was then replaced with fresh DI water for another time period of at least 4 hours and this process was repeated for a total of 6 dialysis cycles. The dialyzed product, an aqueous solution of PHA, was then frozen at -80°C and lyophilized at ∼-80°C and ∼1.0 Pa (Labconco). Proton nuclear magnetic resonance (^1^H NMR) analysis was used to confirm the degree of pentenoate -ene group substitution (DoS) on the hyaluronic acid backbone after lyophilization.

**Figure 8.**
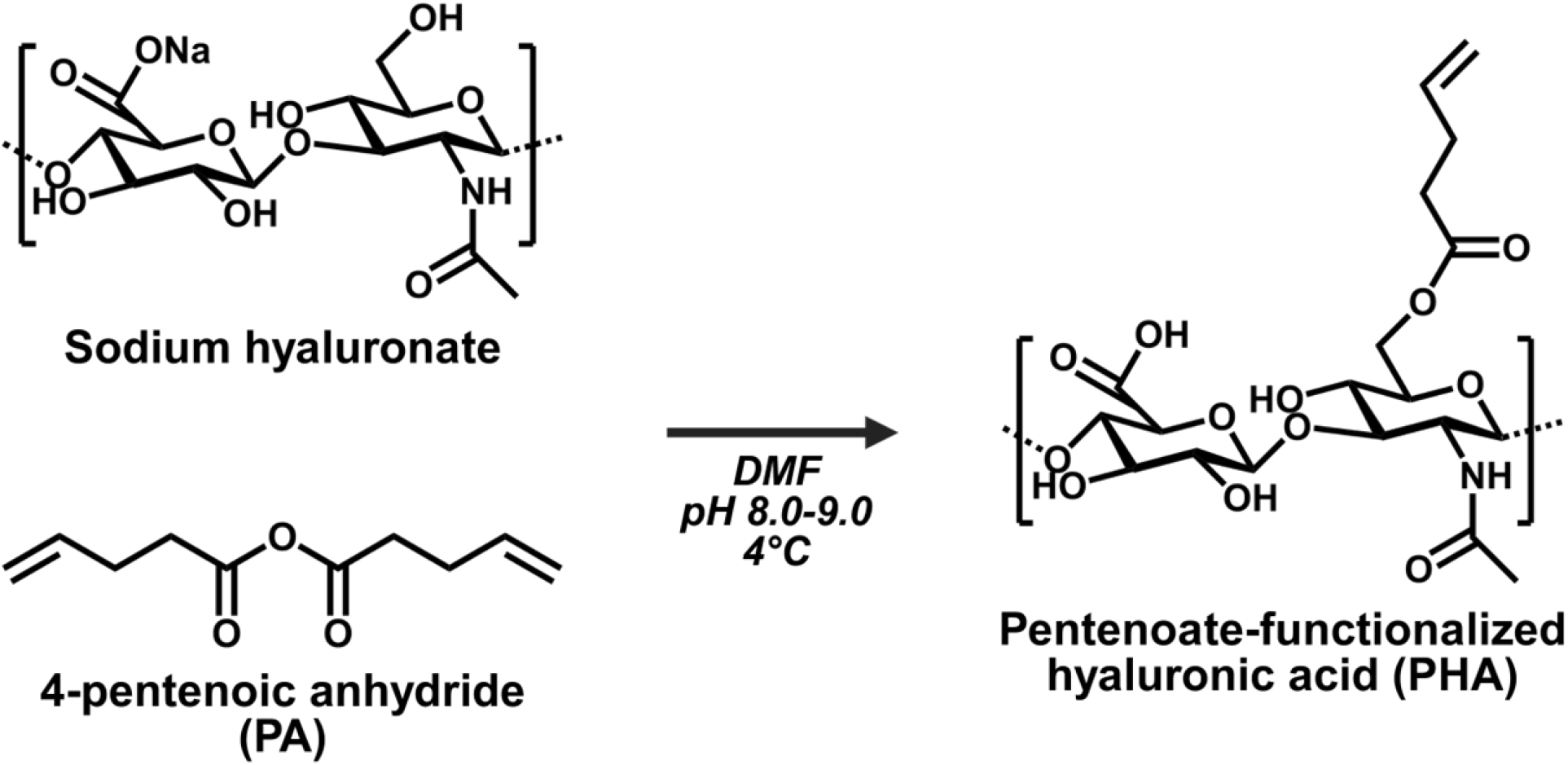
Reaction scheme for the esterification of hyaluronic acid and pentenoic anhydride to yield pentenoate-functionalized hyaluronic acid (made using ChemDraw and BioRender).

### 6.2. Hydrogel synthesis and crosslinking parameters

Hydrogel precursor solutions were formed by combining lyophilized ∼22% DoS PHA, dithiol poly(ethylene glycol) (PEG-DT) (Creative PEGWorks, MW 1 kDa), and lithium phenyl-2,4,6-trimethylbenzoylphosphinate (LAP) (Millipore Sigma) in PBS to achieve a final concentration of 1 mM LAP, the desired PHA weight percentage (w/v), and the desired molar ratio of PEG-DT thiol groups to PHA -ene groups. After all components were combined, the hydrogel precursor solutions were mixed manually, vortexed, and then centrifuged to remove air bubbles. For all studies except photorheology experiments and cell encapsulation experiments, the hydrogel precursor solutions were pipetted into a mold comprised of a silicone sheet (McMaster-Carr, 1.25 mm thickness) between two rectangular glass plates (borosilicate without UV protection). The molds were secured with binder clips and then irradiated with UV light (365 nm UV wavelength, 6 mW/cm^2^ UV intensity) for 60 s on each side (Excelitas OmniCure Elite S2000). **Figure 9** shows the complete crosslinking scheme.

**Figure 9.**
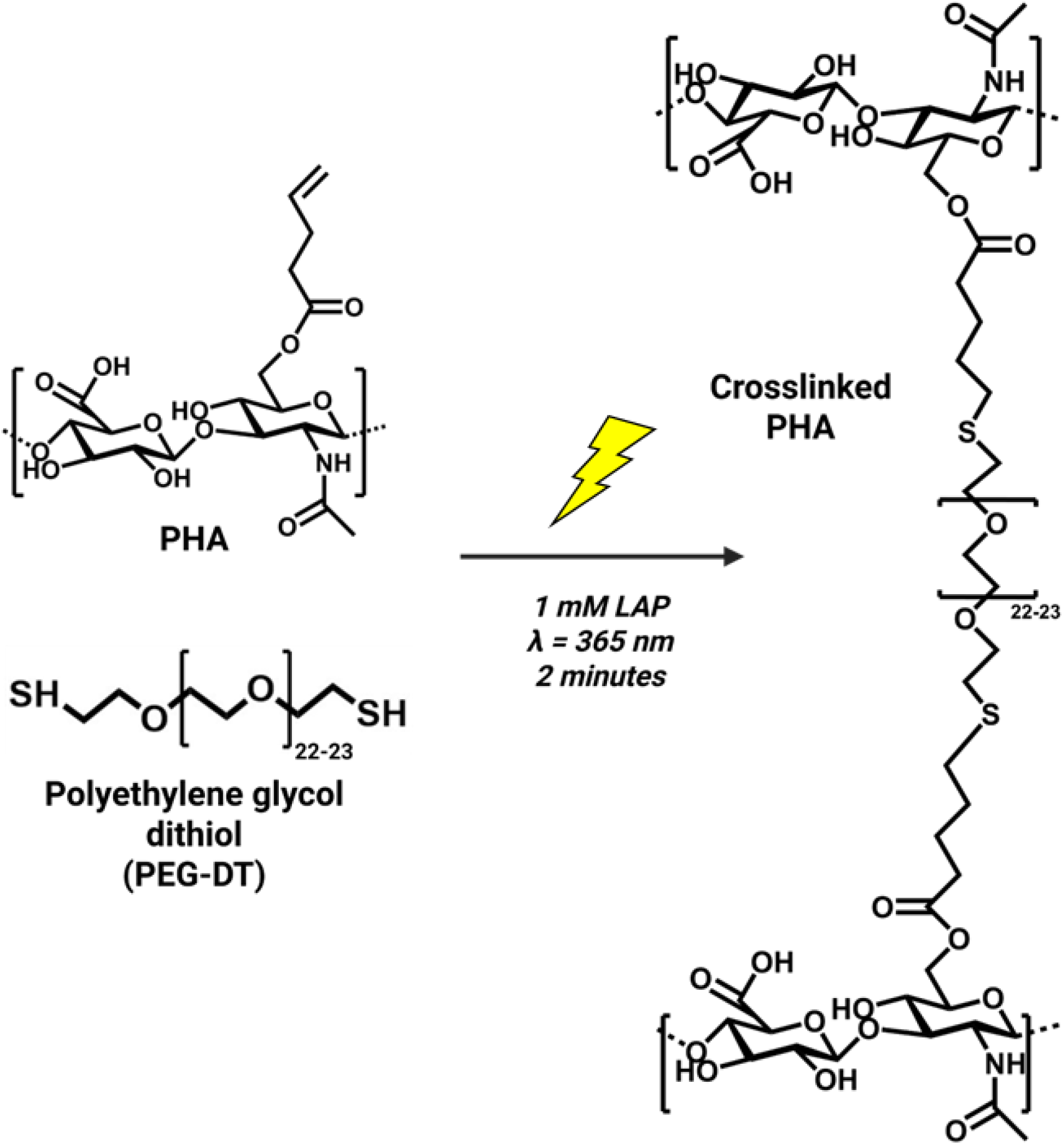
Crosslinking scheme for the photoinitiated thiol-ene reaction between PHA and dithiol polyethylene glycol (PEG-DT) to form a hydrogel network (made using ChemDraw and BioRender).

After irradiation, the hydrogels were removed from the molds, equilibrated in 10 mM NaCl in DI water overnight at 4°C, punched into 8 mm discs, and then equilibrated again in either PBS or 5 µM GFP in PBS for 24 hours at 4°C before use in experiments. **Table 1** summarizes the combinations of polymer concentration and thiol:-ene molar ratio investigated in this study. **Table 2** summarizes all crosslinking parameters used to synthesize hydrogels in this study. Of note, PHA DoS, LAP concentration, UV crosslinking time, UV wavelength, UV intensity, crosslinker identity, and solvent were all kept constant when synthesizing all PHA hydrogels.

**Table 2.**
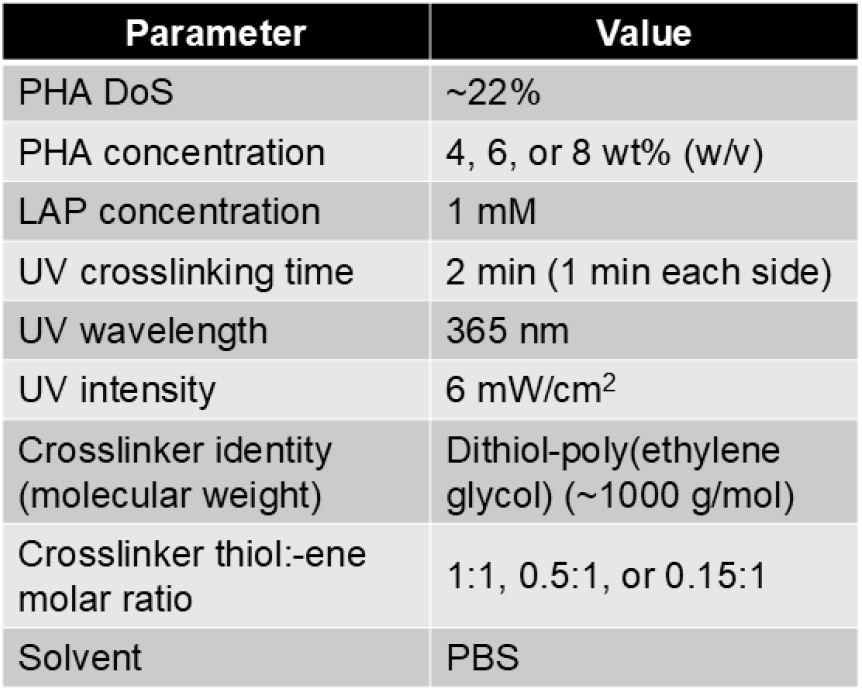
Summary of crosslinking parameters.

### 6.3 Rheological characterization

Rheological characterization was performed using a Discovery Hybrid Rheometer-2 (TA Instruments) equipped with the 8 mm parallel plate geometry (x = 8 mm, y = 8 mm) at room temperature (∼25 °C).

#### 6.3.1. Photorheology

Photorheology experiments were performed to assess the photoinitiated crosslinking kinetics of the PHA hydrogel system. The rheometer was equipped with a 365 nm LED UV-curing accessory and a disposable acrylic plate. Each test was conducted using a constant 0.1% oscillation strain at a constant frequency of 1 Hz with a gap height of 1000 µm. After PHA hydrogel precursor solution was prepared as described above and pipetted onto the plate, a 60 s dwell time elapsed and then the UV light was turned on for 180 s at 6 mW/cm^2^ irradiation intensity. To determine the plateau shear storage modulus for each hydrogel sample tested, the shear storage modulus values (G’) measured during the last 40 seconds of UV irradiation were averaged. At least 3 replicates were measured for each formulation.

#### 6.3.2. Post-swelling strain amplitude sweeps

Strain amplitude sweeps were performed to assess the mechanical properties of the PHA hydrogels after photoinitiated crosslinking and equilibration. The rheometer was equipped with a Peltier plate. PHA hydrogels were synthesized as described above, equilibrated in 10 mM NaCl in DI water overnight at 4°C, punched into 8 mm discs, and then equilibrated again in PBS for 24 hours at 4°C before use in experiments. Each sample was blotted dry with a KimWipe before testing. To minimize slippage, 400 grit sandpaper was affixed to both the lower Peltier plate and the upper plate. The upper plate was lowered onto each sample at a constant 1.0 N axial force until an axial compressive force above 0.03 N was registered, indicating that the equilibrated PHA hydrogel sample was flush with the parallel plate geometry. A strain amplitude sweep was then conducted from 0.01% strain to 100% strain at a constant frequency of 1 Hz. At least 3 replicates were measured for each formulation.

Using the TRIOS software (TA Instruments) the onset point was determined for each sample’s shear storage modulus (G’) curve. The onset point is taken as the intersection of two lines, one drawn tangent to the horizontal portion of the curve, representing the linear viscoelastic region, and one drawn tangent to point at which the magnitude of the slope of the transition region of the curve is maximized, near the curve’s inflection point. This onset point, or yield point, is the limit of the linear viscoelastic region, or the point at which the sample begins to soften or flow. The onset point is unique to the constant frequency at which the oscillation strain sweep was performed (1 Hz). After determining the onset point for each sample, the shear storage modulus of each equilibrated sample was calculated as the average of storage modulus values collected in the linear viscoelastic region of each curve, or the average of storage modulus values corresponding to oscillation strain values less than the oscillation strain value at the onset point.

### 6.4. Molecular Dynamics simulations

The thiol-ene crosslinking reaction between PHA and PEG-DT was modeled using Molecular Dynamics (MD) simulations. Large-scale Atomic/Molecular Massively Parallel Simulator (LAMMPS) was used.^30^ The interatomic potentials were described by the Polymer Consistent Force Field (PCFF).^31^ Each simulation box contained 10 PHA chains with 30 monomers per chain. Six of the 30 monomers were randomly chosen to be functionalized with pentenoate -ene functional groups to simulate a DoS of 20%. Three different crosslinker molar ratios (0.5, 0.77, and 1 thiol groups to reactive -ene groups) were investigated by adding 15, 23, and 20 PEG-DT molecules, each consisting of 4 monomers, to the simulation box. The molecular weights of the PHA polymer and the PEG-DT crosslinker molecules used in the simulations were approximately 20% of the experimental values. Reducing the simulation in terms of molecule size and number provides a higher density of crosslinking sites within the simulation box volume, which ensures a sufficient number of crosslinking events occur within the typical MD simulation time. Initial configurations and force field parameter assignments were generated using the Enhanced Monte Carlo package.^32^

All simulations were first minimized and equilibrated for 0.5 ns using the NVT ensemble at 300 K. An additional 2.5 ns of equilibration was then performed under the NPT ensemble, which includes heating the system from 300 K to 600 K followed by cooling the system back to 300 K. A timestep of 0.5 fs was used throughout the equilibration process. Subsequently, thiol-ene crosslinking reactions were modeled using the *fix bond/react* algorithm^33^ implemented in LAMMPS. This algorithm identifies potential reaction sites by matching topologies of the -ene and thiol groups to a pre-reaction structure template (**Figure 10**). A crosslinking reaction occurs once the terminal carbon atom in an -ene group approaches the sulfur atom in a thiol group within the cutoff distance of 4.5 Å. When this occurs, the topologies of the -ene and thiol groups will change to a post-reaction structure (**Figure 10**). The crosslinking simulations are performed for 0.5 ns using a timestep of 1 fs. Three replicate simulations were conducted for each crosslinking molar ratio investigated and the reported crosslinking efficiency values are the average of crosslinking efficiency values from those three replicate simulations..

**Figure 10.**
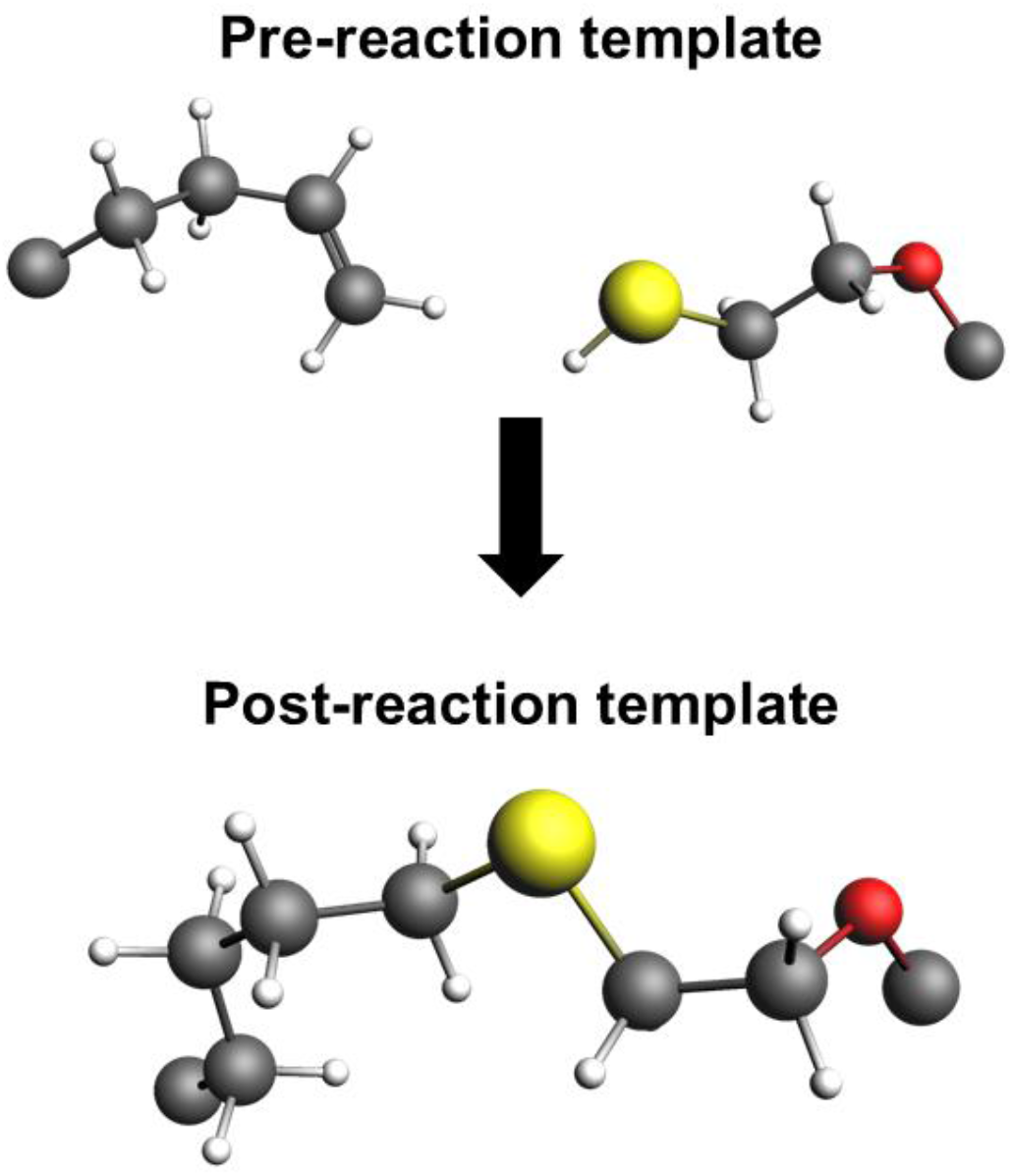
The pre- and post-reaction templates used for Molecular Dynamics simulations. White represents hydrogen atoms, grey represents carbon atoms, red represents oxygen atoms, and yellow represents sulfur atoms. Pre-reaction template represents an unreacted PHA with functional -ene group (left side) and an unreacted dithiol molecule (right side) where the sulfur atom is represented in yellow. Post-reaction template represents a complete thiol-ene reaction. Data visualization provided by Yiwen Zheng.

### 6.5. Fluorescence recovery after photobleaching

Fluorescence recovery after photobleaching (FRAP) experiments were used to measure the diffusion of GFP through the PHA hydrogels to investigate relative mesh size. The protocol for FRAP experiments was adapted from Francis et al.^34^ PHA hydrogels were synthesized as described above, equilibrated in 10 mM NaCl in DI water overnight at 4°C, punched into 8 mm discs, and then equilibrated again in 5 μM GFP in PBS for 24 hours at 4°C before use in experiments. A Leica SP8X confocal microscope equipped with a 405 nm laser and a 25X water immersion objective was used for FRAP experiments. A scan speed of 0.518 s was used. Two pre-bleach images were taken before bleaching was performed to create bleach spots of approximately 10 μm in diameter. After bleaching, recovery images were taken over the course of ∼130 s (20 images at 0.518 s intervals, 20 images at 1 s intervals, 10 images at 10 s intervals). The normalized intensity of the pixels within the bleach spot in each recovery image and corresponding recovery time can be fit to a simplified diffusion equation using modified Bessel functions. This equation and the definition of *τ*_*D*_ (the characteristic diffusion time) are given in Equation 1 and Equation 2, respectively.

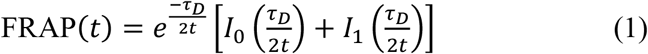

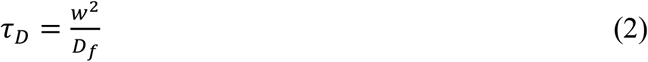

In these equations, FRAP(*t*) is the fluorescence of the bleach spot at a given recovery time (*t*) after bleaching, *w* is the radius of the bleach spot, and *D*_*f*_ is the diffusion coefficient of GFP through each sample. After fitting the normalized intensity values and the corresponding recovery times to Equation 1 and solving for *τ*_*D*_, Equation 2 was used to solve for *D*_*f*_. For each hydrogel formulation investigated, 3 hydrogel samples were tested and 6 sets of FRAP images (from 6 unique bleach points) were obtained from each hydrogel sample.

### 6.6. Cell encapsulation and metabolic activity analysis

For this tunable PHA hydrogel system to be used as a feasible three-dimensional *in vitro* model of meniscus tissue, the encapsulation of meniscal cells inside the hydrogel must be demonstrated as well as the ability of meniscal cells to thrive in 3D cell culture after encapsulation. To encapsulate cells inside the PHA hydrogel system for 3D culture, PHA hydrogel precursor solutions were prepared as described above except for the addition of primary human meniscal cells (isolated from 59-year-old female meniscal tissue resected during total knee arthroplasty (passage 4) according to IRB Exemption #STUDY00018647, NWBioTrust, Seattle, WA) to the hydrogel precursor solution before UV irradiation. Specifically, a 2x PHA solution was prepared in PBS and a second solution containing a 2x concentration of PEG-DT and a 2x concentration of LAP (2 mM) in PBS was also prepared. Cells were lifted, counted, pelleted, and then the cell pellet was resuspended in the second solution containing PEG-DT and LAP. Then, both solutions were combined, mixed manually, and pipetted into a mold for UV irradiation. After irradiation, the hydrogels were removed from the molds, immediately punched into 6 mm discs, and equilibrated in sterile cell culture media (DMEM +10% FBS +1% P/S) overnight at 37°C. See **Figure 11** for a summary of this process.

**Figure 11.**
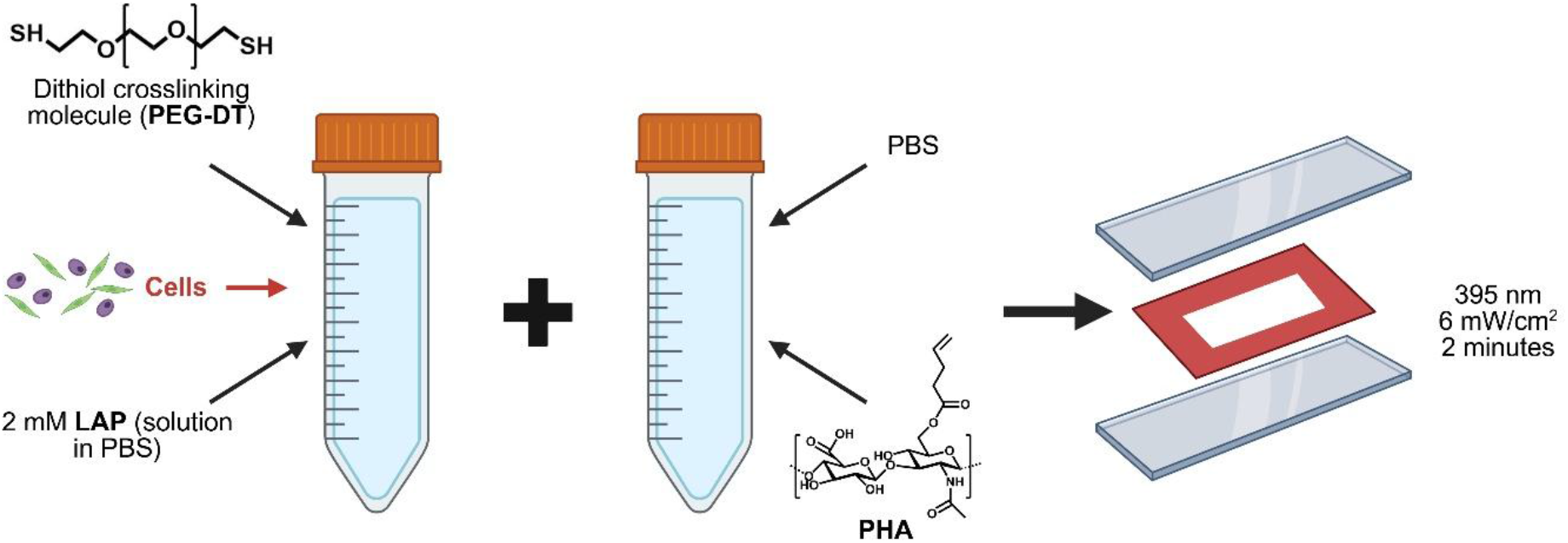
Schematic summarizing cell encapsulation into the PHA hydrogel system for 3D cell culture (made using BioRender).

To assess metabolic activity of encapsulated cells, a PrestoBlue assay was used. PrestoBlue (ThermoFisher Scientific) measures cell metabolism via the measurement of the oxidation of NADH to NAD+. The PrestoBlue reagent contains resazurin which is reduced to resofurin by live cells in culture and this reduction is accompanied by the oxidation of NADH to NAD+. Resofurin is fluorescent (560 nm excitation/590 nm emission), so fluorescence intensity after incubation of cells with the PrestoBlue reagent can be used as a proxy for metabolic activity. The PrestoBlue assay was performed according to the manufacturer protocol. Briefly, the PrestoBlue reagent was added to wells containing PHA hydrogels with encapsulated cells to reach a concentration of 10% PrestoBlue reagent in cell culture media. Samples were incubated for 60 min at 37°C, and then the fluorescence of each well was read using a fluorescent plate reader (Tecan Life Sciences). The PrestoBlue reagent was then removed and wells were washed with fresh cell culture media and then culture continued. This assay was performed every day for 7 days. Each experimental group contained at least 3 replicates.

### 6.7. Statistical analysis

All experiments were conducted with n ≥ 3 samples. For each data set, the data normality was assessed using a Shapiro-Wilk test in GraphPad Prism. For analyses in which all data was determined to be normally distributed, a one-way Brown-Forsythe and Welch ANOVA test was run (assuming unequal variances). For analyses in which not all data was determined to be normally distributed, a nonparametric Kruskal-Wallis test was run. To determine significant differences between hydrogel pre-equilibrium and post-equilibrium shear storage moduli (**Figure 3**) and between metabolic activity levels of cells encapsulated in hydrogels of different formulations at different culture time points (**Figure 7**), a two-way ANOVA test was employed. Significance level was set to 0.05 and asterisks were used to indicate levels of significance: * indicates p≤0.05, ** indicates p≤0.01, *** indicates p≤0.001, and **** indicates p≤0.0001.

